# Novel functional insights from the *Plasmodium falciparum* sporozoite-specific proteome by probabilistic integration of 26 studies

**DOI:** 10.1101/2020.06.18.158899

**Authors:** Lisette Meerstein-Kessel, Jeron Venhuizen, Daniel Garza, Emma J. Vos, Joshua M. Obiero, Philip L. Felgner, Robert W. Sauerwein, Marynthe Peters, Annie S.P. Yang, Martijn A. Huynen

**Affiliations:** Center for Molecular and Biomolecular Informatics, Radboud University Medical Center, Nijmegen, The Netherlands; Radboud Center for Infectious Diseases, Medical Microbiology, Radboud University Medical Center, Nijmegen, The Netherlands; Department of Physiology and Biophysics, School of Medicine, University of California Irvine, Irvine, CA, USA

## Abstract

*Plasmodium* species, the causative agent of malaria, have a complex life cycle involving two hosts. The sporozoite life stage is characterized by an extended phase in the mosquito salivary glands followed by free movement and rapid invasion of hepatocytes in the human host. This transmission stage has been the subject of many transcriptomics and proteomics studies and is also targeted by the most advanced malaria vaccine. We applied Bayesian data integration to determine which proteins are not only present in sporozoites but are also specific to that stage. Transcriptomic and proteomic *Plasmodium* data sets from 26 studies were weighted for how representative they are for sporozoites, based on a carefully assembled gold standard for *Plasmodium falciparum (Pf)* proteins known to be present or absent during the sporozoite life stage. Of 5418 *Pf* genes for which expression data were available at the RNA level or at the protein level, 1105 were identified as enriched in sporozoites and 90 specific to them. We show that *Pf* sporozoites are enriched for proteins involved in type II fatty acid synthesis in the apicoplast and GPI anchor synthesis, but otherwise appear metabolically relatively inactive, in the salivary glands of mosquitos. Newly annotated hypothetical sporozoite-specific and sporozoite-enriched proteins highlight sporozoite specific functions. They include PF3D7_0104100 that we identified to be homologous to the prominin family, which in human has been related to a quiescent state of cancer cells. We document high levels of genetic variability for sporozoite proteins, specifically for sporozoite-specific proteins that elicit antibodies in the human host. Nevertheless, we can identify nine relatively well-conserved sporozoite proteins that elicit antibodies and that together can serve as markers for previous exposure.

Our understanding of sporozoite biology benefits from identifying key pathways that are enriched during this life stage. This work can guide studies of molecular mechanisms underlying sporozoite biology and potential well-conserved targets for marker and drug development.

**Author Summary:** When a person is bitten by an infectious malaria mosquito, sporozoites are injected into the skin with mosquito saliva. These sporozoites then travel to the liver, invade hepatocytes and multiply before the onset of the symptom-causing blood stage of malaria. By integrating published data, we contrast sporozoite protein expression with other life stages to filter out the unique features of sporozoites that help us understand this stage. We used a “guideline” that we derived from the literature on individual proteins so that we knew which proteins should be present or absent at the sporozoite stage, allowing us to weigh 26 data sets for their relevance to sporozoites. Among the newly discovered sporozoite-specific genes are candidates for fatty acid synthesis while others might play a role keeping the sporozoites in an inactive state in the mosquito salivary glands. Furthermore, we show that most sporozoite-specific proteins are genetically more variable than non-sporozoite proteins. We identify a set of conserved sporozoite proteins against which antibodies can serve as markers of recent exposure to sporozoites or that can serve as vaccine candidates. Our predictions of sporozoite-specific proteins and the assignment of previously unknown functions give new insights into the biology of this life stage.

## Introduction

Malaria is a mosquito transmittable disease resulting in over 220 million clinical cases and half a million deaths annually. Most deaths are caused by *Plasmodium falciparum (Pf)*, one of the five species of *Plasmodium* that can infect humans. The infection begins with the deposition of liver-infective sporozoite forms in the skin by blood-feeding mosquitoes. These sporozoites travel to the liver where they invade, differentiate and multiply asymptomatically inside hepatocytes for approximately a week before releasing red blood cell (RBC)-infective merozoites into the circulation. The subsequent asexual multiplication, rupture and re-invasion of the parasites into circulating RBCs cause the symptoms associated with malaria.

Identifying specific sporozoite proteins will aid in the understanding the biology of this highly motile stage in which the parasite is directly exposed to the body’s immune system, similar to blood stage infective merozoites. Sporozoites deposited in the skin can take up to 90 minutes to reach the liver [1], a much longer potential exposure to immune system than the 90 seconds of merozoites. Mosquito midgut-derived and circulating sporozoites are less infectious for hepatocytes than those that have resided in the salivary gland [2, 3]. The ability to improve and remain infectivity in the salivary gland of mosquitoes for an extended period of time (approximately 1 week) is an intriguing phenomenon that is poorly understood [4]. It is suggestive of a molecular landscape where the parasite is kept in low activity but can quickly activated to evade immune systems, invade and develop in hepatocytes. Understanding the specific molecular make-up of sporozoites will shed light on this aspect of its biology and may reveal new targets for interventions.

A number of studies have identified genes that are expressed in sporozoites during both their development in mosquito midgut (oocyst) and in the salivary gland. They have identified expression at the RNA level[5] and at the protein level[6] with a specific emphasis on surface proteins[7]. The studies varied in their focus and resolution, with RNA level studies facing the challenge of their relevance for protein expression[8] while proteomics studies face challenges in detecting low abundance proteins. More importantly, while such studies address the question what is present in the sporozoite, they do not address what is specific to it. Therefore, to prioritize sporozoite specific proteins, we implemented a naïve Bayesian data integration in which both transcriptomics and proteomic data were included. For informed data integration, gold-standard lists representing proteins that are either sporozoite specific or non-sporozoite specific were manually assembled from the existing literature on individual proteins. Using this gold standard, published *Plasmodium* datasets were weighted by the presence of sporozoite specific proteins and the absence of proteins known to be absent from sporozoites, allowing us to obtain an extended list of predicted sporozoite specific proteins. We classified 90 proteins as sporozoite-specific, of which 67 were not part of the gold standard. “Conserved, hypothetical proteins with unknown functions” were examined using sensitive homology and orthology detection tools [9, 10] to predict their function and shed light on the biology of sporozoites.

From the proteins that in the assembled proteomics data were identified to be present in sporozoites, we examined whether we can identify a limited set of proteins for which human antibodies are generated[11] and that can serve as or markers of exposure to sporozoites. As attractive targets would be conserved between different *P. falciparum* strains, we sequenced three *P. falciparum* strains, NF54, NF135 and NF166, and required selected proteins to have limited genetic diversity between those strains as well as in sequenced genomes in PlasmoDB[12].

## Methods

### Sporozoite protein and transcriptomic data sets

The data integration was performed using transcriptomic and proteomic data sets from 22 studies describing all life cycle stages of *P. falciparum* (**S1 Table**). Data sets were obtained from PlasmoDB version 43 [12], and literature (**S1 Table**). In addition to that, transcriptomics data on the liver and blood stage of *Plasmodium cynomolgi* and the sporozoites of *Plasmodium vivax* were included, as well as proteomics studies covering the sporozoite stage of *P. vivax* and the liver stage of *P. yoelii*, respectively, resulting in a total of 48 data samples from 26 different studies (**S1 Table**). Data from non-*falciparum* studies were converted into *P. falciparum* IDs using the orthologs lists available on PlasmoDB, favouring orthologs that are syntenic in the case of multiple options.

### Gold standards

Bayesian data integration requires gold standards, in this case of proteins known to be highly enriched in sporozoites or depleted from them. We required positive gold standard proteins to be present dominantly in sporozoites based on western blot or immunofluorescent assay data, resulting in a selection of 31 sporozoite-specific proteins (**S2 Table**).

The negative gold standard was curated by searching literature for non-sporozoite proteins. We selected 19 proteins based on their literary evidence of absence from sporozoites. Additionally, 20 gametocyte specific proteins [13] were added to the negative gold standard, increasing the total number to 39 negative gold standard proteins (**S3 Table**). These proteins spanned the remaining *P. falciparum* life cycle stages in the human host, with the majority being found in (a)sexual blood stages.

### Bayesian data integration

The used data sets were examined for their correlations with each other (**S1 Figure**). By and large most data sets show little correlation. We did leave the few correlated data sets in to keep the data integration transparent and maximize the amount of included information. Oocyst-derived and salivary gland sporozoites showed high correlations with each other, which led us to combining all respective studies into the “sporozoite” data input and not make a distinction between those stages.

Proteomic data were converted into unique peptide counts for each protein identified and transcriptomic data were converted into expression percentiles for a total of 5668 *P. falciparum* gene IDs. Proteomic and transcriptomic data was binned consistently for all data sets, with 0, 1, or >1 identified unique peptides, or into four bins, containing transcripts that are in the >80 percentile, >60 percentile, >40 percentile and <40 percentile, respectively. The data sets were then weighted according to their ability to retrieve the gold standard proteins. Each bin in each data set was given a log2 score according to equation (1), where B = present in bin, S = sporozoite specific and nonS = not sporozoite specific.

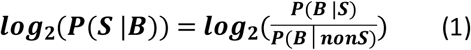

The Bayesian score for an individual protein is then the sum of the scores for the bins in which it occurs (one bin per data set). As the expected number of sporozoite-specific proteins was unknown, no *prior* was included, rather we used cutoffs based on the position of known sporozoite specific proteins to define sporozoite specific proteins and sporozoite enriched proteins.

### Overrepresentation of function categories in sporozoite proteins

GO terms were acquired from PlasmoDB and formatted into a .gmt file according to the format specified by the GSEA server at the Broad Institute[14]. GSEAPreranked on the ranked list of sporozoite-specific proteins was used with the conservative “preranked” option in the “classic mode”, i.e. using Kolmogorov Smirnoff statistics to determine enriched GO terms. The large variable protein families RIFIN, STEVOR and PfEMP1 (*var* genes) were left out of the analysis. A new GO term for gliding motility in *Plasmodium* was assembled by searching literature for proteins associated with gliding motility in sporozoites, ookinetes and merozoites. For this we extended the list of glideosome associated proteins assembled by Lindner *et al.*[15] with 10 new proteins possibly associated with the glideosome. These proteins were either annotated with the motility GO term GO:0071976 (SSP3, CelTOS, LIMP protein, plasmepsin VII and glideosome-associated connector), or otherwise known to be involved in gliding motility (CTRP[16], SIAP1[17], GAPDH[18], GAP40[19], IMC1l[20]). Similarly we added a list of GPI-anchored proteins to the set of processes for which we examined enrichment using the list of Gilson *et al.*[21], based on the hypothesis that the type II fatty acid synthesis was required for the creation of GPI anchors [22]. To investigate Pfam domain enrichment in sporozoite proteins, Pfam annotations for all proteins were downloaded from PlasmoDB and were used as gene sets in the GSEA. To prevent the GSEA analysis results for enriched pathways to be affected by the large numbers of hypothetical proteins in *P. falciparum*, those were filtered out before the analysis. Furthermore, proteins that were part of the gold standard were left out of the analysis to prevent circular arguments. Proteins for which no transcriptomic and proteomic data were available were left out as well. Manual examination of the Gene Set Enrichment Analysis results showed that some gene sets were significantly enriched in sporozoites only because they were depleted from the low scoring proteins. Therefore, we required for the significantly enriched processes in sporozoites that they actually contained proteins in the set of “sporozoite enriched proteins.”

### Sequencing of *P. falciparum* strains

NF54 originates from West Africa, and is isolated from a woman infected in the Netherlands nearby an airfield[23]. The NF135 clone is a clinical isolate that originates from Cambodia[24]. The NF166 clone is a clinical isolate from a child that visited Guinea[25]. Whole genome sequencing of the three strains was performed with Illumina NextSeq 500, resulting in raw paired-end fastq reads of 151 base pairs (bp).

### Quality control and trimming

To examine the quality of the raw fastq reads, FastQC (version 0.11.5) was used[26]. The Nextera Transposase sequence contamination at the 3’ ends of the reads were trimmed off with a stringency of 12 bp, using Trim Galore (version 0.4.3)[27]. CleanNextSeq_paired was used to remove excess of G’s from 3’ ends of the reads after 100 bp[28]. Only reads with a minimal length of 35 base pairs were retained.

### Alignment and variant calling

The *P. falciparum* 3D7 reference genome (v3.0, PlasmoDB, plasmodb.org, 14 chromosomes, mitochondrial genome and apicoplast genome) was indexed and the trimmed fastq reads were aligned to the reference genome using Bowtie2 (version 2.2.8) with the local alignment setting[29]. The SAM files obtained were converted to BAM files and subsequently sorted and indexed with SAMtools (version 1.4.1)[30]. SNPs and indels were called using SAMtools and BCFtools (version 1.4.1). Alignments were visually inspected with the Integrative Genomics Viewer (IGV, version 2.3.98)[30]. Coverage was calculated with BEDtools (version 2.26.0)[31].

### Filtering of SNPs and indels

Filtering of SNPs and indels was performed with BCFtools (version 1.4.1)[30]. SNPs with a base Phred quality (Q) > 30 were used for further analysis. Furthermore, we required that the proportion of high quality bases (the DP4 scores in the VCF files) supporting the indel or the SNP in >= 75% of the called bases to include them for further analysis.

### Effect of SNPs and indels on protein sequences

The effect of the mutations on the predicted protein sequences was determined using the Genomic Ranges package in R [32]. Amino acid sequences of all proteins were compared to amino acid sequences of the reference genome and with each other. Any variation at the amino acid level between reference and the examined strain and among the examined strains was denoted.

### Homology detection

Homology detection of *P. falciparum* proteins with unknown function was done using HHPred[9] with default settings (HHblits, 3 iterations). For orthology detection we used best bidirectional best hits at the level of sequence profiles[10].

### Detecting a set of proteins that elicit antibodies in all CPS volunteers

We used a greedy “set cover” algorithm that in each step selects, from a list of evolutionary conserved sporozoite proteins (less than eight nonsynonymous SNPs per kb in PlasmoDB), the one that is immunogenic (elicits antibodies) in the highest numbers of volunteers for which no immunogenic protein was already in the set. In the analysis from Obiero *et al.* [11], some long proteins were split into separate peptides. Those were analyzed separately by the algorithm: i.e. as if they were separate proteins. When multiple proteins were immunogenic in the same number of volunteers for which no immunogenic protein had been selected yet, we chose from those the one that had the highest immunogenicity in all the volunteers. We furthermore required genes to have maximally two non-synonymous SNPs in the combined NF135 and NF166 strains.

## Results

### Data integration

At least 26 studies, containing 48 data sets have been published of *P. falciparum, P. cynomolgi*, *P. vivax* and *P. yoelii* transcripts and proteins at various developmental stages, including 11 that were specific to the sporozoite stage (**S1 Table**). In order to optimally exploit those data to obtain sporozoite enriched proteins we integrated them in a Bayesian manner (Methods). Integrating the data sets using the sets of 31 positive and 39 negative gold standard proteins (**S2 Table, S3 Table**) produced a list of all proteins in *P. falciparum* ranked according to their likelihood of being sporozoite-specific (**S4 Table**). The score distribution of the negative and positive gold standard proteins varied depending on using all available data sets, or proteomic or transcriptomic data separately (Figure 1A and **S2 Figure**). The Bayesian integration using only transcriptomic data sets resulted in a ranked list where 14 negative gold standard genes scored higher than the lowest scoring positive gold standard gene (**S2 Figure**). This overlap was lower when using only proteomic data sets (**S2 Figure**), but still contained 8 proteins. The least overlap between positive and negative gold standard members was observed when combining transcriptomic and proteomic data (Figure 1A). This also produced an outlier, STARP. Although this protein was identified as being a sporozoite protein [33], in our combined data sets it scored 42 times lower than the second to last scoring positive gold standard protein. It therewith does not appear specific to sporozoites in the available data. STARP was excluded from the score distribution of gold standard proteins that was used to define the sporozoite specific proteins (see below), but gold standard list and integration where not changed *post hoc*. Based on the observed overlaps, we decided to continue our research using the ranked list based on the combined proteomic and transcriptomic data sets.

**Fig 1.**
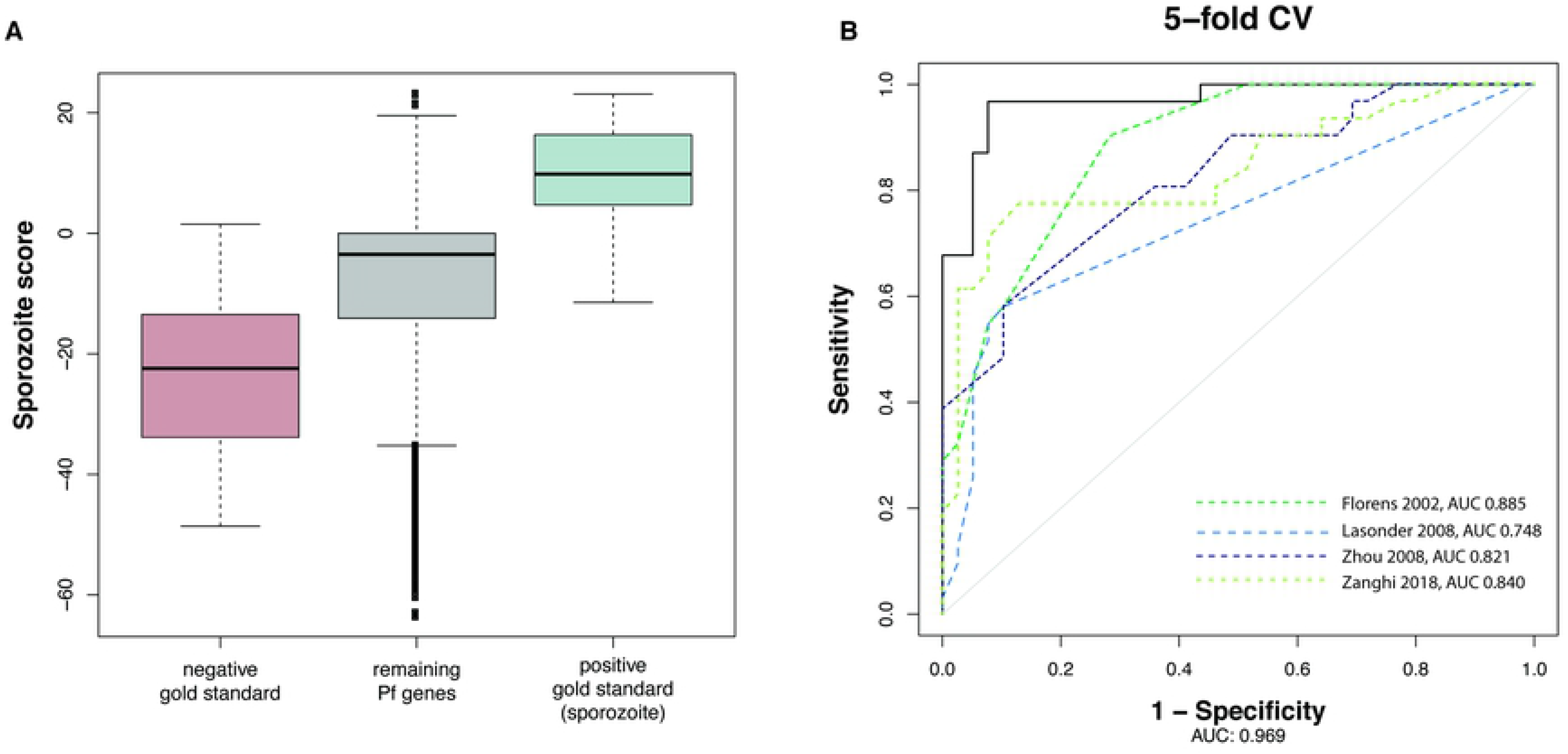
Bayesian data integration identifies sporozoite-specific genes in *P. falciparum*. **A)** Bayesian score distributions of the proteins from the negative gold standard, the positive gold standard and the remaining proteins, **B)** cross validation of our predictions (5-fold cross validation) with all data sets (Black solid line) and predictions with individual data sets (dashed coloured lines).

### Sporozoite specific proteins and sporozoite enriched proteins

Proteins were considered *sporozoite-specific* when ranking in the first quartile of the gold standard protein list i.e. scoring above 12.27, which represents a factor of being at least 2^12.37^ to 2^~ 5200^ more likely to be specific to sporozoites than to any of the other stages. Do note that hereby we ignore the *prior* probability of a protein being sporozoite specific at all. Although such a prior is standard in Bayesian data integration, in this case a prior, e.g. of 1/5, is given the high cut-off for sporozoite specific proteins that we used, not very relevant. As we did not consider STARP to be sporozoite specific (see above), we considered proteins that scored higher than the second to lowest positive gold standard protein (LIMP protein, score = −1.31), to be *enriched* in sporozoites. Finally, the abundance of unique peptides by mass-spectrometry was assessed for each protein. Proteins were deemed *present* in sporozoites when identified in two independent studies or with more than 1 unique peptide in at least one study. Our analysis thus identified 90 sporozoite-specific proteins, 1105 sporozoite-enriched proteins and 2736 that were present in sporozoites (**S4 Table**). Out of the 90 sporozoite-specific proteins, 67 were not part of the positive gold standard list. We validated our predictions by 5-fold cross validation (5-fold CV) by randomly skipping 1/5^th^ of the gold standard proteins from the data integration and assessing their predicted sporozoite specificity based on the remaining data (Fig 1B). The high sensitivity and specificity indicated that novel sporozoite-specific proteins would also score higher than non-sporozoite specific proteins. We also compared the ranking of the gold standard proteins based on the integrated data with a ranking based on individual data sets of sporozoite RNAs and proteins. The cross validation separated the gold standard proteins better than individual data sets, supporting the integration of multiple data sets (Fig 1B).

### Function prediction of non-annotated proteins

Many of the genes in the *P. falciparum* genome encode hypothetical proteins with unknown molecular function [34]. The fraction of unknowns that is specific for sporozoites (32 out of 90, 36%) is at least as high as in the rest of the genome (32%). To improve understanding of their potential functions, we examined the proteins for domains with known functions from any species using sensitive homology detection with HHpred[9], combined with manual examination of conserved residues. We compared sporozoite specific proteins with PFAM domains, the human proteome, and the proteome of *Toxoplasma gondii*, an apicomplexan related to *P. falciparum* that is present in the HHpred database. The sporozoite specific protein with the highest score that was not part of the gold standard, PF3D7_0104100, showed (barely) significant sequence similarity with the prominin family (E=10E-4) and low levels of sequence identity with e.g. human prominin-2 (12%). To cross-check the homology of PF3D7_0104100 with prominin we examined homology with *T. gondii* proteins. The sequence similarity between the *P. falciparum* protein and *T. gondii* TGME49_218910 (E=3.4e-44) and between the *T.gondii* protein and the human prominin-2 were highly significant (E=2.3e-21), indicating that PF3D7_0104100 is indeed member of the prominin family. PF3D7_0104100 has like the prominin family five (predicted) transmembrane regions with most of the protein localized outside the cell (Figure 2). Analysis of a sequence alignment with orthologs in other *Plasmodium* species reveals the conservation of ten cysteine residues that are all predicted to be extracellular (Figure 2). Such extracellular cysteines can form disulphide bonds as has been observed for other extracellular *Plasmodium* protein domains [35] and has been suggested for human prominin[36]. Two pairs of the extracellular cysteines were conserved between the human prominin and PF3D7_0104100 (Figure 2, **S3 Figure**). We were able to predict molecular functions of three other sporozoite specific proteins and 27 sporozoite enriched proteins using HHpred and best bidirectional hits with human proteins at the level of sequence profiles (**S6 Table**)[10].

**Fig 2.**
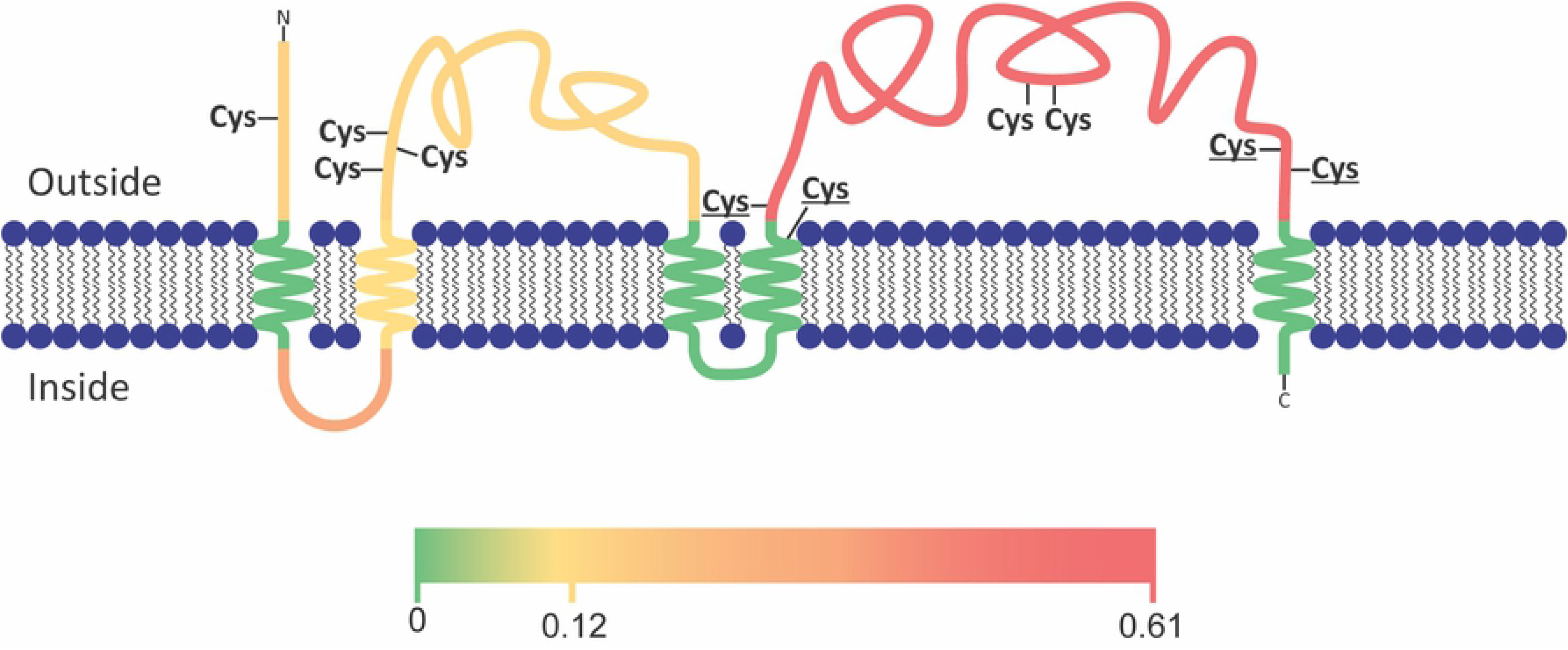
Predicted membrane topology of Pf3D7_0104100, a sporozoite-specific protein that is homologous to the prominin/CD133 protein family. The level of polymorphisms among *P. falciparum* strains is indicated for the separate regions as the average number of polymorphisms per nucleotide in the strains in PlasmoDB. PF3D7_0104100 has a high density of polymorphisms within *P. falciparum* strains that are concentrated in the second extracellular loop. Cysteines that are conserved among the homologs in *Plasmodium* species are indicated. The cysteines that are conserved also in human homologs are in bold. Note that the conserved cysteines occur in close proximity to each other, suggesting the formation of disulphide bonds.

**Fig 3.**
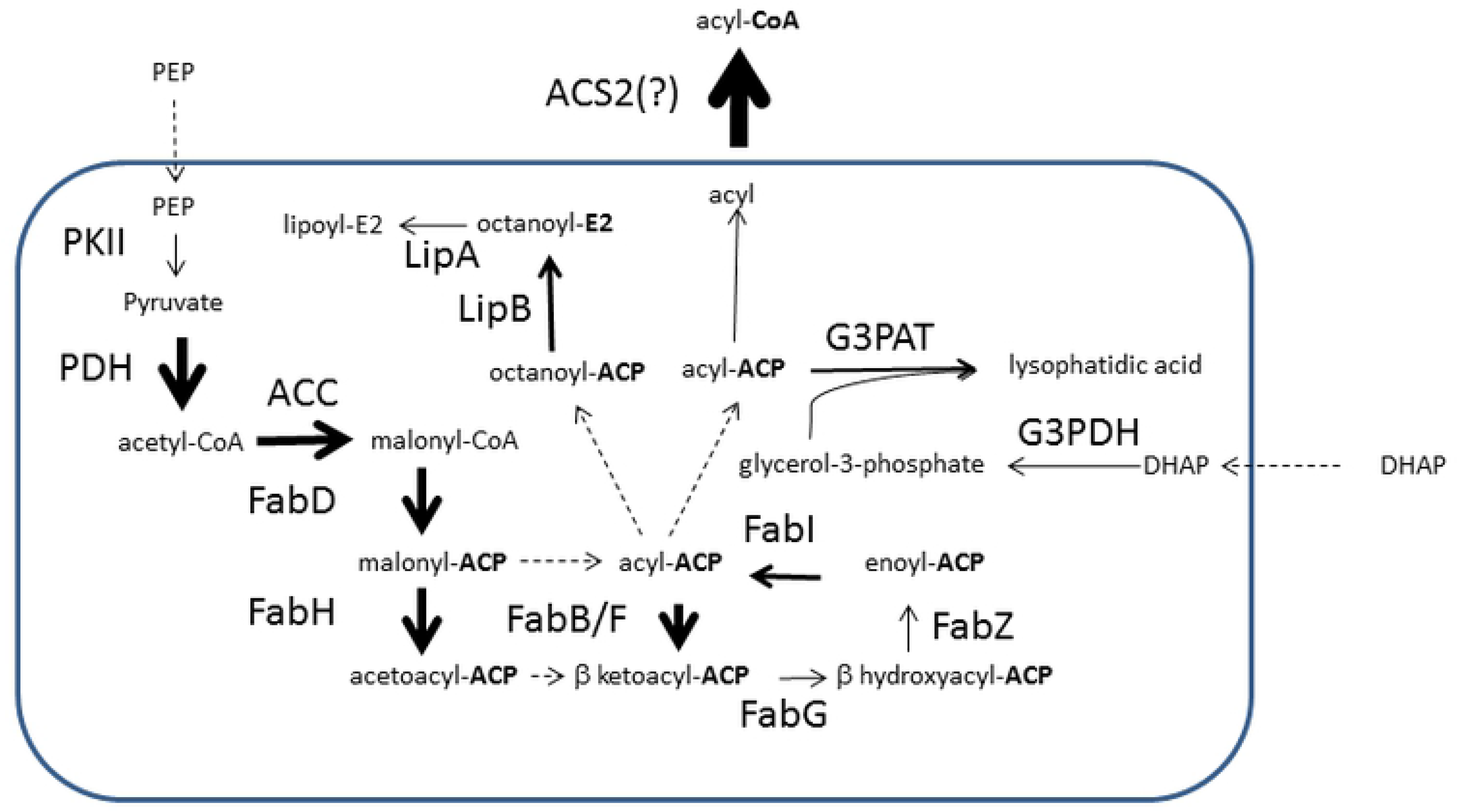
Apicoplast fatty acid synthesis proteins are enriched among sporozoites. The width of the arrows is determined by the Bayesian score reflecting the level of over representation of that enzyme in sporozoites, e.g. 16 for ACS2 and 8 for FabB/F (**S4 Table**). For PDH that consists of three proteins, the width of the arrow was determined by the average of those three. Most of FASII proteins are enriched in sporozoites, except PKII, FABZ and LipA. The scheme is a simplification of the pathway as depicted by Shears *et al.*[22], to which ACS2 was added as it is highly enriched in sporozoites and relevant for fatty acid synthesis.

### Over-represented domains and pathways

We examined whether specific protein domains and pathways were over- or under-represented in the sporozoite proteins using Gene Set Enrichment Analysis[14]. For this we augmented the list of proteins involved in specific processes from PlasmoDB with the glideosome and glycosylphosphatidylinositol (GPI) anchor proteins that we considered specifically relevant to sporozoites (Methods)[14]. At the domain levels there was an over-representation of three protein domains: PF09175 (also named Plasmod_dom_1, unknown function), PF08373 (the RAP domain, putatively RNA binding) and PF12879 (the C-terminal domain of the SICA proteins that are associated with parasitic virulence). Two of these three domains are either unique to *Plasmodium* species (PF12879) or to *P. falciparum* (PF09715). At the level of pathways, we observe significant enrichment of the Type-II fatty acid synthesis (FAS-II) and the GPI anchor biosynthesis pathways. The enzymes present in the FAS-II pathway are located in the apicoplast – a reduced plastid-like organelle that shares similarity with chloroplasts of algae and plants [37]. This over-representation is consistent with its essentiality for sporozoite development [38]. Interestingly, *P. falciparum* has fourteen Acyl CoA synthetases [39], of which the distribution over the various fatty acid metabolizing processes is largely unresolved. There is one Acyl CoA synthetase, ACS2, that is specific to sporozoites and that would be an interesting candidate to convert the fatty acids produced in the apicoplast to acyl-CoA. Curiously, there was no enrichment in sporozoites of members of the acyl-ACP thioesterase family (PF3D7_1135400 and PF3D7_0217900). Acyl-ACP thioesterases play an important step in the pathway by hydrolyzing the acyl moiety from the ACP before it can be converted to acyl-CoA.

The second enriched pathway is GPI-anchor biosynthesis, with three proteins (GPI mannosyltransferases 1, 2 and 3: PF3D7_1210900, PF1247300 and PF3D7_1341600) enriched in sporozoites. GPI represents a class of glycolipids found as either free lipids or attached to proteins [40, 41]. Surface parasite proteins (such as the merozoite surface proteins 1, 2 and 4) anchor to the parasite cell-membrane via GPI moieties [42]. Sporozoites shed surface proteins (such as circumsporozoite protein and thrombospondin related anonymous protein) when moving [43]. With GPI being used to anchor proteins to the parasite surface, it would have to be synthesized constantly to replace the surface proteins shed during motility. There was no enrichment of the glideosome or of GPI anchor proteins among proteins specific to the sporozoite. However, 39 of the 52 glideosome associated proteins are “present” in the sporozoite stage (75%), representing a slight overrepresentation when compared to the 2,736 of 5,447 proteins (50,2%) detected in sporozoites (Fisher’s exact, P=0.0018). Similarly, there are 19 out of 28 GPI-anchor proteins identified in Gilson *et al.*[21] among the proteins present at the sporozoite stage, but they are not specifically enriched in sporozoites and hence are also expressed in other developmental stages. The fact that proteins involved in movement (such as glideosome and GPI) are enriched but not specific to sporozoites indicated that sporozoite share a common movement machinery with other the life-cycle stages.

### Protein functions of sporozoite-specific proteins in the context of sporozoite biology

Apart from examining over represented pathways we also examined the individual sporozoite specific proteins with regard to their potential function in sporozoite biology. One of the interesting features of sporozoites is their inactivity in the sporozoite glands, which is among others maintained by translational repression with the Puf2 protein [44]. In view of the observation that the translational repression occurs in two waves [44], it is interesting that the sporozoite specific protein PF3D7_0411400 contains a Dead box helicase like its paralog of DOZI that is involved in translational repression in gametocytes of *Plasmodium berghei* [45]. From a gene expression regulatory perspective are furthermore interesting: three Zinc finger proteins (PF3D7_0615600, PF3D7_0521300, PF3D7_1008600), and the transcription factor AP2-04 that has mainly been implicated in ookinetes[46]. With respect to the epigenetic regulation, the histone deacetylase sir2b (PF3D7_1451400) that is involved in epigenetic silencing of var gene expression[47] is sporozoite specific, in contrast to its paralog sir2a that is not enriched in sporozoites. Examining the sporozoite specific signaling proteins with respect to their potential function in maintaining the temporal inactivity of the parasite we noted the RAC-beta serine/threonine protein kinase PfAKT (PF3D7_1246900) and two 14-3-3 domain proteins (PF3D7_1422900 and PF3D7_1362100). The PfAKT that is involved in Artemisinine resistance [48] is specifically interesting in view of the role of its metazoan orthologs, AKT1/2/3, in regulating cellular metabolism to support cell survival [49]. In human it phosphorylates a large number of targets, a number of which are after phosphorylation bound by 14-3-3 proteins. Two 14-3-3 proteins are sporozoite specific: Pf14-3-3I (PF3D7_1422900) and PF3D7_1362100. Pf14-3-3I has been shown to bind phosphorylated Histone H3 [50]. The Raf kinase inhibitor RKIP (PF3D7_1219700), which may be a substrate of the calcium-dependent protein kinase 1 [51], may also be involved in keeping sporozoites in an inactive state as its human ortholog in inhibits the MAPK pathway [52]. Furthermore, with respect to inactivity of sporozoites it is interesting to mention that Prominin-1/CD133, one of the two human orthologs of the Prominin gene (PF3D7_0104100), is known as a marker for dormant stem cells, e.g. in melanoma [53] or in Neural cells [54], and has been shown to activate the PI3K/AKT pathway[55]. Together, these data suggest that these proteins may also modulate sporozoite survival via a mammalian Akt-like pathway as previously shown in asexual blood stages.

Finally, the presence of the tubulin polymerization promoting protein p25 alpha (PF3D7_1236600) as well as thioredoxin-like protein (PF3D7_0919300) and thioredoxin-like associated protein 1 (PF3D7_1230100) whose orthologs are associated with tubulin in *T. gondii* [56] are worth noting in view of the essential role of the microtubules in the sporozoite stage [57]. In contrast to these proteins, the β-tubulin, the two α-tubulins and the other thioredoxin-like associated proteins associated with tubulin [56] are not enriched in sporozoites. Given the presence of β- and α-tubulins in all stages of the life-cycle, PF3D7_1236600, PF3D7_0919300 and PF3D7_1230100 may be crucial players involved in providing sporozoites with their characteristic, thin cylindrical shape.

### Under-represented pathways

In contrast to the paucity of upregulated processes, there is significant under representation (FDR =< 0.001) of processes linked to splicing, translation, translation elongation, folding of proteins as well as proteolysis (S5 Table). These processes are primarily modulated by a number of sporozoite specific proteins involved in transcriptional silencing e.g. PF3D7_0411400 that contains a Dead box helicase domain[45] and PF3D7_1451400, a histone deacetylase involved in the epigenetic silencing of virulence (var) gene expression [47]. Furthermore, there is significant depletion of proteins involved in carbon metabolism: glycolysis and citric acid cycle (S5 Table).

### Selecting sporozoite proteins as targets or markers of past infection

Antibodies that bind sporozoite proteins have been induced in 38 volunteers after chloroquine chemoprophylaxis with *P. falciparum* sporozoites (CPS)[11], Table S7. Induced antibody profiles represent a blue print of immunogenic proteins in sporozoites, liver stages and early blood stages. Here we focus on the set of sporozoite target proteins that may be used as markers of previous sporozoite exposure or may act as potential targets for vaccines. Minimal sequence variation between *Pf* strains would thereby strengthen the candidature of the proteins for epidemiological or clinical applications, however antibody eliciting proteins including CSP, show relatively high levels of polymorphisms among sequenced malaria strains (**S4 Figure**), and also sporozoite specific proteins show high levels of polymorphisms (**S5 Figure**). The selection of proteins that could serve as markers of exposure or vaccine candidates is a compromise between on the one hand protein sequence conservation and on the other hand the frequency of volunteers with antibodies to that protein. As the maximum level of sequence variation between candidate marker proteins we used 8 non-synonymous SNPs per kilobase, which is lower than the variation in for instance Pfs48/45, a highly conserved gametocyte protein with 8.9 nonsynomymous SNP per kilobase. As the minimum number of people in which a protein should elicit antibodies we chose six (out of the 38). Using a “greedy search algorithm” (methods) we selected a set of nine proteins of which at least one elicited antibodies per volunteer (Table 1).

**Table 1:** putative markers of exposure to sporozoites. Genes selected by the greedy method to cover all volunteers with their gene ID, function and number of volunteers with antibodies after CPS immunization as reported by Obiero *et al.* [11]. The non-synonymous SNPs are given as a proxy for genetic variability, with PlasmoDB and two sequenced laboratory strains as reference.

| Gene ID | function | No. (%) of volunteers with antibodies | Non-syn SNP/kb PlasmoDB | Non-syn SNPs |  |
| --- | --- | --- | --- | --- | --- |
|  |  |  |  | NF135 | NF166 |
| PF3D7_0630600 | Conserved hyp | 20 (52.6) | 2.8 | 0 | 1 |
| PF3D7_0906500 | Arginase | 19 (50.0) | 6.5 | 1 | 1 |
| PF3D7_1456700 | Conserved hyp. | 19 (50.0) | 2.9 | 0 | 0 |
| PF3D7_0719700 | 40S ribosomal S10 | 15 (39.5) | 0 | 0 | 0 |
| PF3D7_1219100 | Clathrin heavy chain | 12 (31.6) | 5.5 | 0 | 2 |
| PF3D7_0301700 | Hypothetical exp. | 10 (26.3) | 6.3 | 0 | 1 |
| PF3D7_1122700 | Conserved hyp. | 9 (23.7) | 7.8 | 0 | 0 |
| PF3D7_1455800 | LCCL prot. | 6 (15.8) | 4.1 | 0 | 0 |

### Variation in sporozoite proteins among selected *Pf* strains NF135 and NF166

Aside from the criteria that the proteins selected as markers for sporozoite exposure are conserved in sequenced *P. falciparum* strains in PlasmoDB, we also required them to be conserved in two strains that have been used in research into heterologous protection after CPS: NF135 and NF166. We therefore sequenced these strains, as well as NF54 (Methods). The mean coverage of mapped reads in the coding regions ranged from 28 for NF54 to 44 for NF166 (**S8 Table**). A Phred quality score cut-off of 30 was applied similar to other studies involving *Plasmodium vivax* and *P. falciparum* sequencing and variant calling, [58–61]. Manual examination of SNPs still uncovered SNPs with a high Phred score (> 100) that were polymorphic within a single, in principle haploid genome, possibly reflecting mapping issues in duplicated regions. Next to a Phred score of 30, we also introduced a required the presence of a variant nucleotide in at least 75% of the high quality bases. Both criteria were also applied to the indels. Numbers of called SNPs and indels were roughly similar for NF135 and NF166, reflecting their independent geographic origins. As expected, NF54 showed much lower numbers of called SNPs and indels than the other strains, as it is the parental line from which 3D7 is derived (S8 Table). As observed in Plasmodb, proteins in NF166 and NF135 showed high levels of polymorphisms for antibody-binding proteins enriched in sporozoites (Figure S6). Nevertheless, most of them had few or no variations in the set of proteins that we selected as markers for previous exposure (Table 1). Lowering the maximum allowed variation in these two strains further, e.g. to zero polymorphisms for all proteins, did not allow us to select a set of proteins that together elicited antibodies in all 38 volunteers.

## Discussion

The sporozoites stage of the *Plasmodium* life cycle represents the parasite’s first interaction with the human immune system and can be used to effectively vaccinate against infection [11]. The set of genes and proteins expressed at this stage has been determined in at least 11 studies on *Plasmodium spp*, of which nine on *P. falciparum*, creating a conundrum of which dataset to use when studying sporozoite biology, and when deciding to use multiple datasets, how to integrate the datasets. The individual sporozoite studies did not focus on determining which expression patterns are specific to the sporozoite stage, rather they examined the presence of transcripts or proteins. Such data of course can be integrated by combining them in a relatively straightforward manner as has e.g. been done for sporozoite RNA expression[62], but that does not address how specific the expression of a gene is to sporozoites. By combining RNA and protein expression data measured across life stages with sets of proteins known to be present or absent from sporozoites in a Bayesian manner we have created a single list of proteins ranked by their overrepresentation in sporozoites relative to other stages. Despite translational repression, which is expected to reduce the correlation between mRNA and protein levels, including the mRNA levels led to a better performance at the protein level that only including proteomics data. Cross validation shows furthermore that the integrated list is better at separating the gold standard positive and negative data sets from each other than the individual data sets.

In this study, we did not separate datasets derived from oocyst sporozoites (the earliest form of this stage) and salivary gland sporozoites (the mature form). There may have been subtle difference in protein expression between sporozoites in the two differing host environments (midgut versus salivary gland) that were not detected. However, oocyst-sporozoite data represent a minority of the combined data set (2/26) and are highly correlated with salivary gland-derived sporozoite data (**S1 Figure**). A list of proteins with its own sporozoite specificity score will be a valuable resource for studying sporozoite biology and understanding novel protein function. Genetic manipulations of malaria parasites can only occur during blood stage development, which makes studying proteins that are essential for both blood and sporozoite stages difficult. Using our list in combination with the recently published *piggyBac* whole genome mutagenesis study [63], will allow researchers to determine the approach required for generating a knockout parasite i.e. whether an inducible (for essential blood stage and low sporozoite specificity score) or straight knockout (for a high sporozoite specificity score) system should be considered.

In our analysis, we found an enrichment in the proteins involved in type II fatty acid synthesis which consistent with literature [38]. An increased output of lipids from this pathway may feed into the production of GPI anchors that are made up of different sugar and lipid components. We did observe a slight enrichment in proteins involved in creating GPI anchors, i.e. the three GPI mannosyltransferases. Although there is currently no established relationship between type II fatty acid pathway and synthesis of GPI anchors[22], it may be interesting to pursue it for this stage of the life-cycle given the importance of CSP – the most abundant protein on sporozoites, that is anchored to the surface via a GPI anchor.

Processes involved in the production, folding and catabolism proteins were under-represented in the sporozoite specific and enriched list. Similarly, genes involved in metabolism such as those part of glycolysis and citric acid cycle were also under-represented, suggestive of sporozoites existing in a low metabolic state. Sporozoites are released in the mosquito’s circulation as early as day 12 post blood meal [3] and make their way to the salivary gland. Nevertheless they are generally less infectious for liver cells until after a period of maturation within the salivary gland[64]. Although the exact nature of this maturation is not known, there is evidence that these parasites remain in a latent state until ejected into the human host[4, 65]. To understand how sporozoites exist in this state of inactivity, we have identified several interesting targets such as an ortholog of AKT1/2/3, two 14-3-3 proteins, an ortholog of the raf kinase inhibitor and an ortholog of the quiescent stem cell marker prominin/CD133.

Sequence variation among *Plasmodium* strains is pervasive [66], and is possibly responsible for the limited heterologous protection after CPS vaccination with NF54 and challenge with NF135 and NF166 [67]. Indeed as we have shown here both sporozoite specific proteins and proteins that elicit antibodies are highly polymorphic, and only a fraction of those are conserved between NF54, NF135 and NF166 (Fig S6, S7). The correlation between immunogenicity and level of sequence conservation suggest that antigenic drift plays a role in the sequence variation. It is not clear whether antigenic drift would also be responsible for high variation among sporozoite specific proteins, as we did not observe a correlation between the Bayesian score of sporozoite specificity and the immunogenicity. Nevertheless, among the large number of sporozoite proteins that elicit antibodies there are still proteins that show limited sequence variation and allow selecting of a small set of proteins that are well conserved and against which antibodies could serve as potential vaccine targets or markers of previous exposure to sporozoites.

In summary, we show a set of previously unidentified sporozoite-specific proteins and assign functions potentially related to the enduring state of inactivity of the salivary gland sporozoite. We further identify sporozoite-directed humoral immune responses and their potential as functional or diagnostic responses that can be elucidated in future studies.

---

**S1 Table: Overview of all datasets used for the Bayesian data integration**

**S2 Table: Positive gold standard members**

Evidence for expression in sporozoites

**S3 Table: Negative gold standard members**

Evidence for expression in other life stages

**S1 Figure: Correlations of all integrated data sets with each other, hierarchically clustered.**

Study code (two letters and year) as in table S1. Samples are coded as sporozoite (sporo), salivary gland sporozoite (SGS), oocyst derived sporozoite (ODS) or oocyst (OOC), liver stage (LS), blood stage (BS, ery, asexual, merozoite, schiz, ring) or other stages (gametocyte/gam, zygote, ookinete). Transcriptomics studies are given a percentile (percent) for each gene they detected and proteomics studies a score for each unique peptide (uniq_pept) per gene.

**S2 Figure: Distribution of sporozoite-specific probability score.**

Score for positive and negative gold standard and all remaining Plasmodium falciparum genes (rest) when using only proteomics data as input (A) or only transcriptomics data (B).

**S4 Table: Ranked genes for their sporozoite-specificity**

**S3 Figure: Alignment of PF3D7_0104100 from *P.falciparum*, TGME49_218910 from Toxoplasma gondii and Prominin-2 from Homo sapiens**. The in PF3D7_0104100 predicted transmembrane regions are underlined, the conserved cysteines are boxed. Do note that the fourth transmembrane region is relatively long. Predictions with other tools than TMHMM [1] like Phobius [2] indicate a shorter TM region, putting the cysteine that is located in that TM region in the extracellular space. The *Toxoplasma* protein was included because its sequence profile has significant sequence similarity against both the human protein profile (E= 2e-20) and the *Plasmodium* protein profile (E= 3.4e-44), while the similarity between the human and the *Plasmodium* protein is less significant (E=0.0001).

**S5 Table: Biological processes enriched (top list) or depleted (bottom list) in the sporozoite**

Enrichment based on the relative absence of the proteins involved at an FDR <= 0.01 and an enrichment score > 0.20. The type II Fatty Acid Synthesis are the genes from Figure 2, of which the gold standard genes were left out in the GSEA analysis. The “translocation of peptides or proteins into hosts” GO category did not have any proteins among the sporozoite enriched proteins, and was left out of the description of the results. (N)ES: (normalized) enrichment score, FDR: false discovery rate

**S6 Table: Sporozoite enriched proteins annotated as “unknown function” in PlasmoDB for which orthology with human proteins could be detected using best bidirectional hits at the level of sequence profiles**

**S4 Figure: Number of people in whom antibodies against *Plasmodium* protein were detected by protein microarray after CPS immunization**

Antibody prevalece does not correlate with the number of Non-synonymous SNPs per kb coding region of the respective gene (PlasmoDB).

**S5 Figure: Combined levels of polymorphisms for strains NF135 and NF166 among stage-specific proteins.**

sporozoite proteins, selected at various levels of stringency, gametocyte proteins, and the remaining proteins. Sporozoite enriched or sporozoite specific proteins show relatively high levels of polymorphisms, while gametocyte proteins are clearly depleted of polymorphisms. Furthermore, antigenic proteins of either stage are enriched relative to non-antigenic proteins

**S7 Table: Antibody responses in volunteers after CPS immunization.**

**S6 Figure: Venn diagram with *Plasmodium falciparum* proteins that elicit antibody responses and have non-synonymous SNPs**

SNPs in NF135 (blue) and NF166 (yellow) compared to the reference in NF54/3D7. The grey shaded area contains proteins without any SNPs, they are hence identical to the 3D7 reference and NF54 strain in both NF135 and NF166. The overlap (light green) shows proteins that have SNPs in both NF135 and NF166, and 27 proteins (dark green) have the exact same SNPs in both strains.

**S8 Table: Levels of polymorphism in NF54, NF135 and NF166 relative of the reference strain 3D7**. Information for all annotated Plasmodium falciparum proteins and the proteins considered immunogenic and sequencing depth.

